## Supplementary figures for "Functional brain network reconfiguration during learning in a dynamic environment"

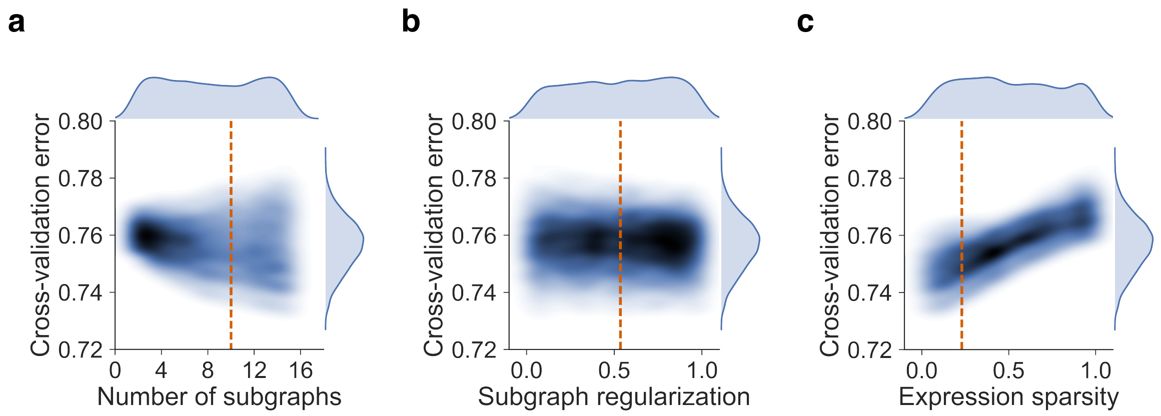


**Supplementary Fig. 1**

Optimal parameters for nonnegative matrix factorization. (a) Number of subgraphs. We randomly sampled the number of subgraphs from a uniform distribution ($k \in [2, 15]$). The contour plot shows the Kernel density of the bivariate distribution. The darker blue area represents the higher probability mass. We selected the optimal parameter ($k = 10$) by averaging the parameter values that ensured that the cross-validation error was in the bottom 25% of the sampling distribution (orange dashed line). (b) Subgraph regularization. We randomly sampled values of subgraph regularization from a uniform distribution ($\alpha \in [0.01, 1.0]$) and select the optimal parameter ($\alpha = 0.535$). (c) Expression sparsity. We randomly sampled values of expression sparsity from a uniform distribution ($\beta \in [0.01, 1.0]$) and select the optimal parameter ($\beta = 0.230$).

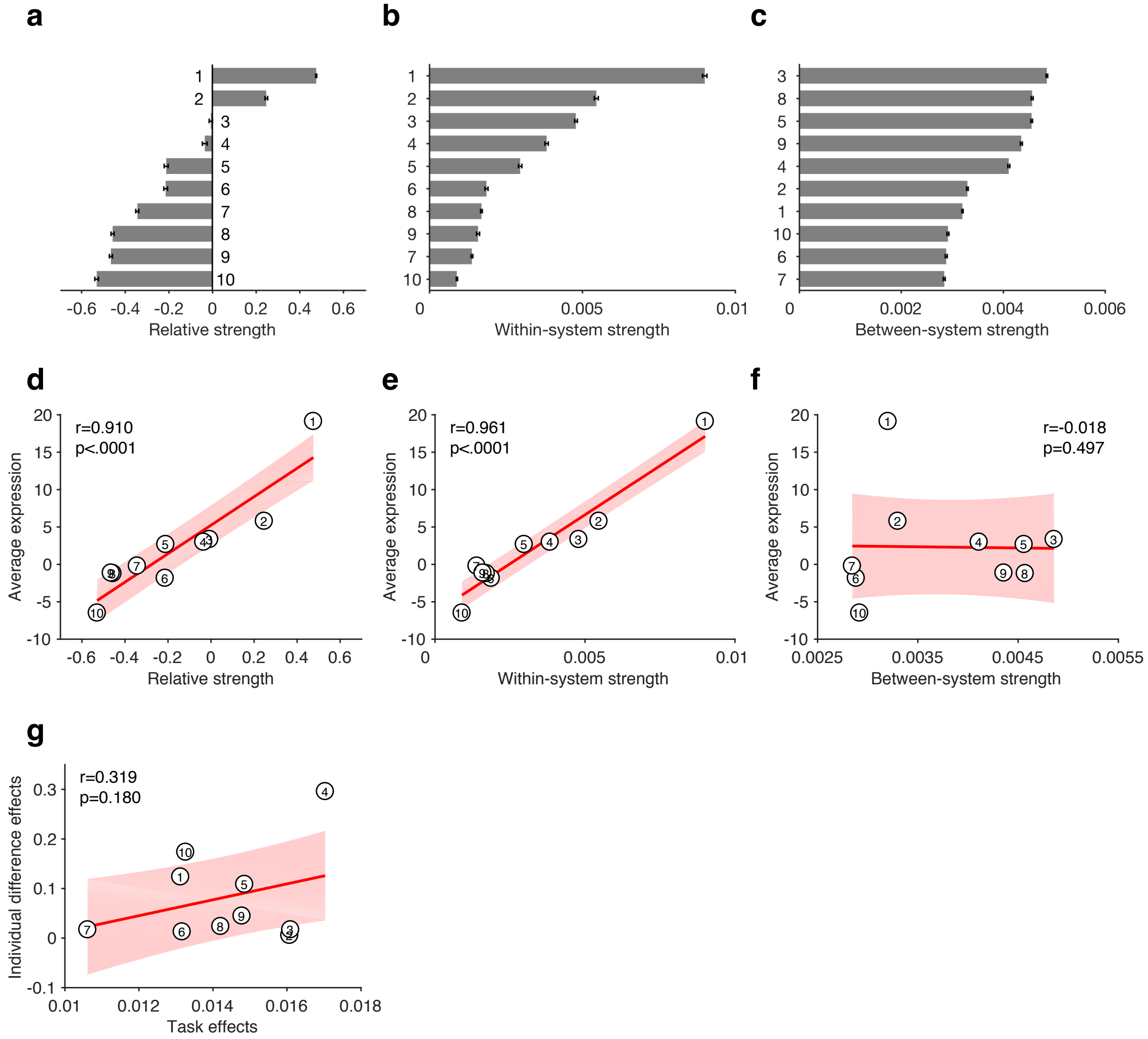


**Supplementary Figure 2**

Properties of subgraphs. (a) Subgraphs differ in the extent of within- versus between-system edge strength. For each subgraph, the strength of within-system edges (edges linking two nodes that both belong to the same system; Fig. 3b) was averaged, and the strength of between-system edges (edges linking one node from one system to another node from another system; Fig. 3b) was averaged. The ten subgraphs are ordered according to the relative strength of within- versus between-system edges. To form a normalized relative strength, we subtracted the average strength of between-system edges from the average strength of within-system edges and then divided this difference by their sum. A high relative strength means that a subgraph has stronger within-system edges than between-system edges (e.g., subgraph 1). The 95% confidence interval of each subgraph was estimated by boostrapping 10,000 times on the edges of that subgraph. (b) Subgraphs differ in the extent of within-system strength. For each subgraph, the strength of within-system edges was averaged. For demonstration, the ten subgraphs are ordered according to within-system strength. The 95% confidence interval of each subgraph was estimated by boostrapping 10,000 times on the edges of that subgraph. (c) Subgraphs differ in the extent of between-system strength. For each subgraph, the strength of between-system edges was averaged. For demonstration, the ten subgraphs are ordered according to the between-system strength. The 95% confidence interval of each subgraph was estimated by boostrapping 10,000 times on the edges of that subgraph. (d) The relationship between relative strength and average expression across subgraphs. Average expression was calculated as the difference between positive expression and negative expression. Each data point represents one subgraph. A significantly positive correlation was observed. The red line represents the regression line and the shaded area represents the 95% confidence interval. (e) The relationship between within-system strength and average expression across subgraphs. A significant positive correlation was observed. The red line represents the regression line and the shaded area represents the 95% confidence interval. (f) The relationship between between-system strength and average expression across subgraphs. There was no significant correlation. The red line represents the regression line and the shaded area represents the 95% confidence interval. (g) The relationship between explained variance of task effects and explained variance of individual difference effects across subgraphs. For the task effects, we implemented a regression model that examined the influence of CPP, RU, reward and residual updating on temporal relative expression for each subgraph. Explained variance was indexed as $R^2$ of the regression model. For the individual difference effects, we implemented a regression model to predict individual normative learning for each subgraph. We included two regressors: dynamic modulation of normative factors (CPP and RU) on subgraph expression, and average subgraph expression. We then calculated $R^2$ of the regression model for each subgraph. Among the ten subgraphs, subgraph 4 showed the strongest $R^2$ for both task effects and individual difference effects.

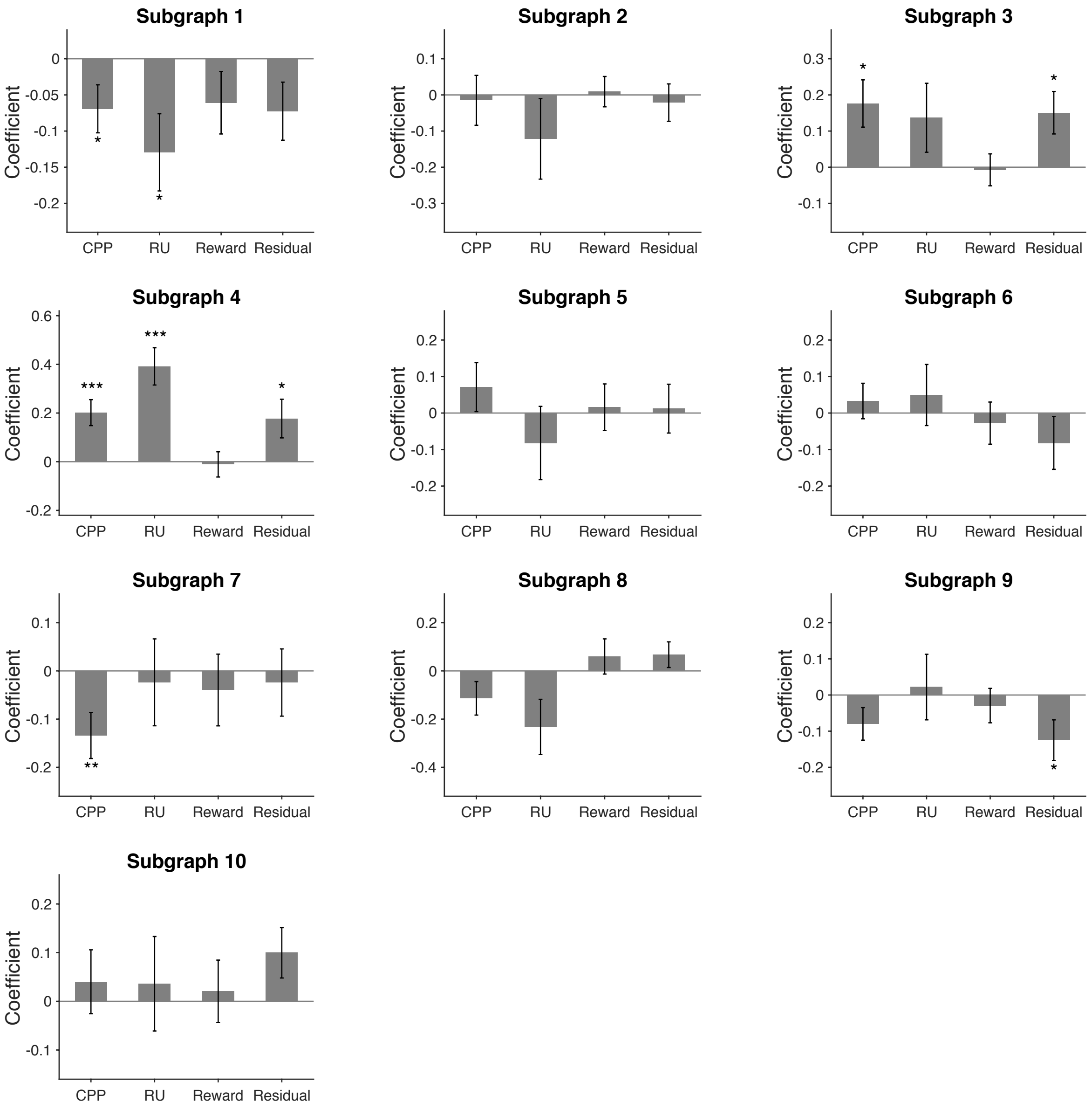


**Supplementary Figure 3**

Modulation of temporal expression by learning factors. A regression model that included CPP, RU, reward and residual updating as predictors of trial-by-trial expression was fitted for each participant and subgraph, and then regression coefficients were tested on the group level. Source data are provided as a Source Data file. Error bars represent one SEM. (*$p$<0.05, **$p$<0.01, ***$p$<0.001)

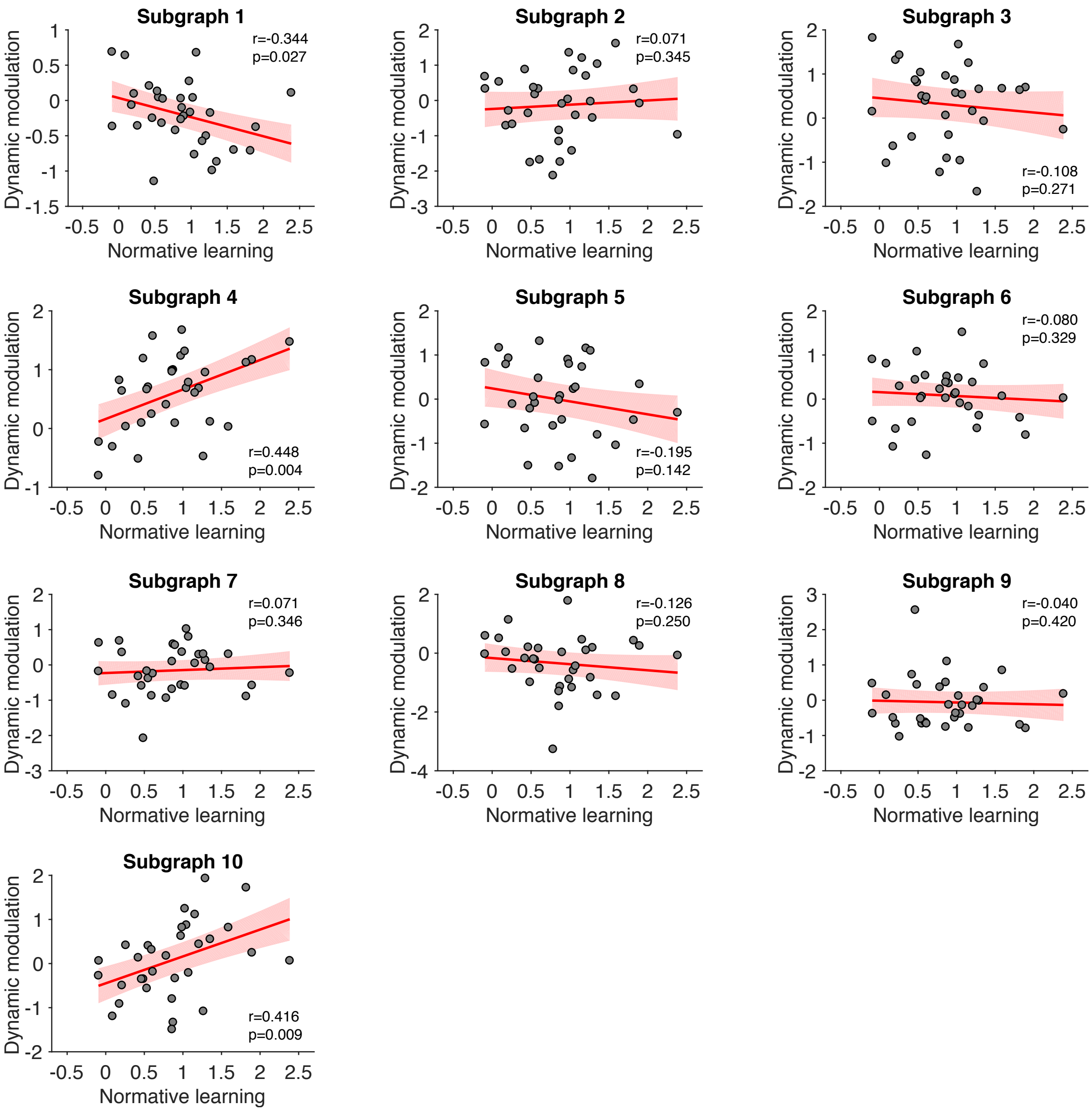


**Supplementary Figure 4**

The relationship between individual normative learning and the dynamic modulation of subgraph expression by normative factors. Normative learning was indexed by the sum of the coefficients of CPP and RU from a behavioral regression model, and represents the extent to which a participant's behavior was influenced by the two normative learning factors. Dynamic modulation was indexed by the sum of coefficients of CPP and RU from the regression model against trial-by-trial expression in Supplementary Fig. 3, and represents the extent to which subgraph expression in that participant was influenced by the two normative learning factors. Each point represents one participant. The red line represents the regression line and the shaded area represents the 95% confidence interval. Source data are provided as a Source Data file.

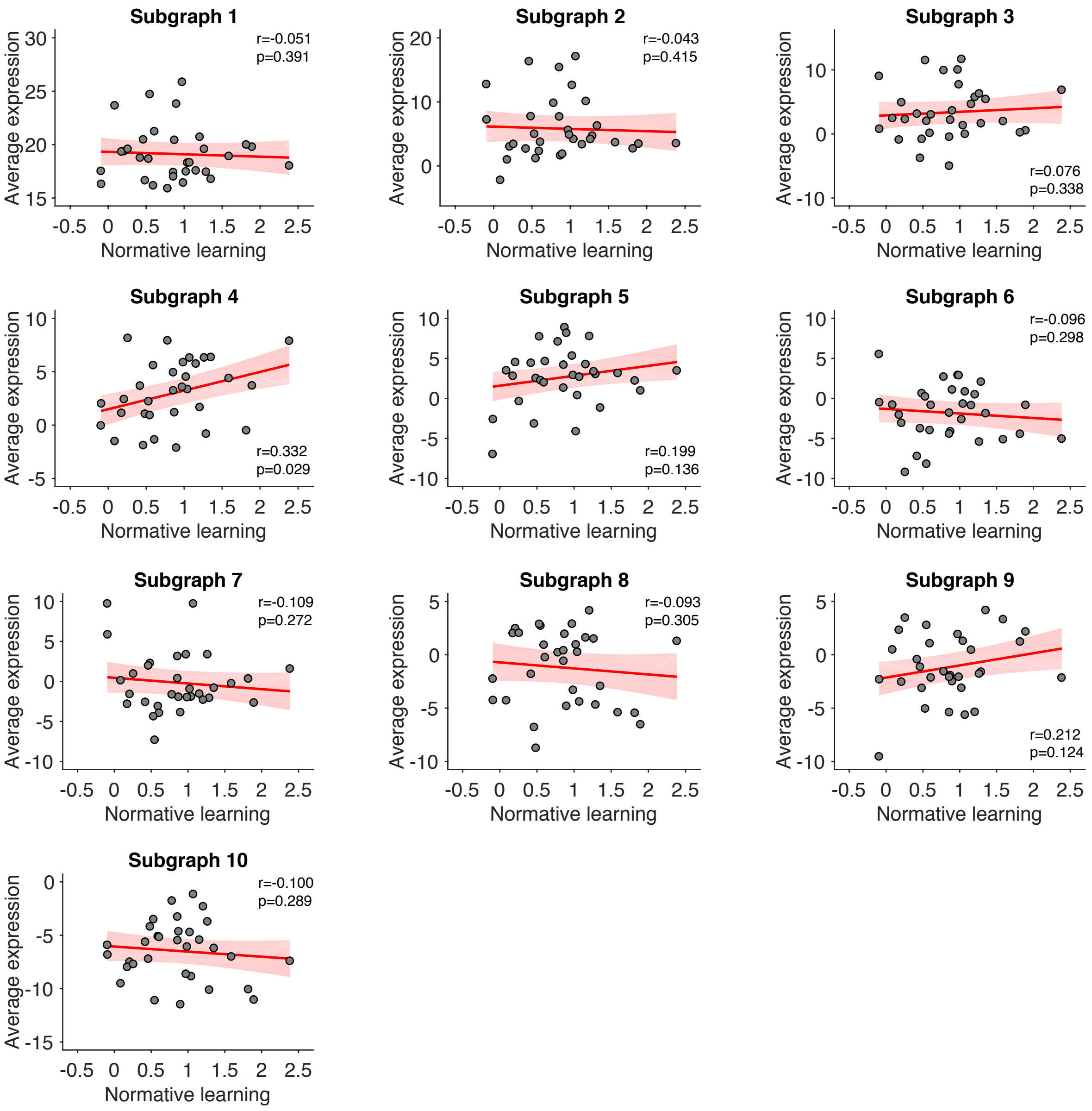


**Supplementary Figure 5**

The relationship between individual normative learning and the average expression of each subgraph. Each point represents one participant. The red line represents the regression line and the shaded area represents the 95% confidence interval. Source data are provided as a Source Data file.

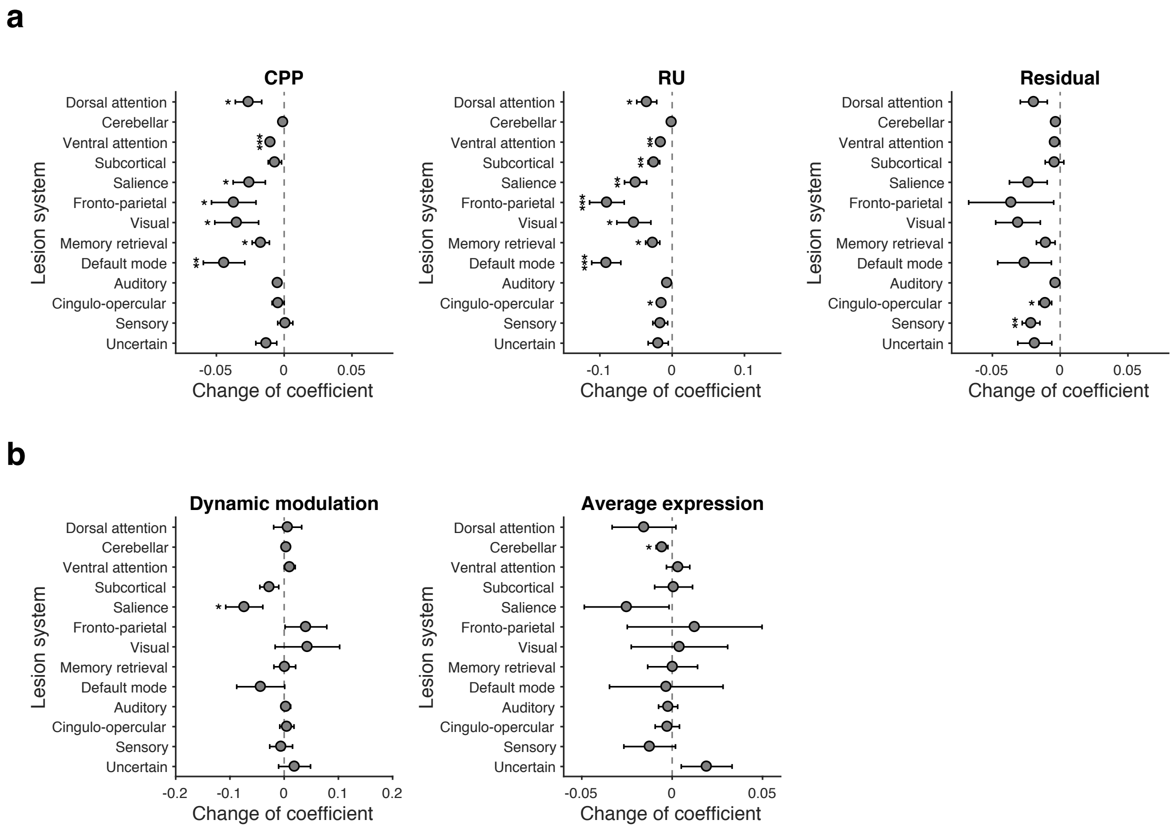


**Supplementary Figure 6**

Contributions of different functional systems in subgraph 4. (a) Contributions of different functional systems to the effect of learning factors on temporal expression of subgraph 4. We removed all edges from one of the 13 systems and re-estimated the coefficients for CPP, RU, reward and residual updating. Then we compared these coefficients with the original coefficients (including all the edges) to estimate the contribution of each system. Error bars represent one SEM. (*$p$<0.05, **$p$<0.01, ***$p$<.001) (b) Contributions of different functional systems to the relationship between normative learning and dynamic modulation and average expression of subgraph 4. We repeated the same procedure and estimated the change of correlation coefficients for each relationship separately. Source data are provided as a Source Data file. Error bars represent one SEM. (*$p$<0.05)

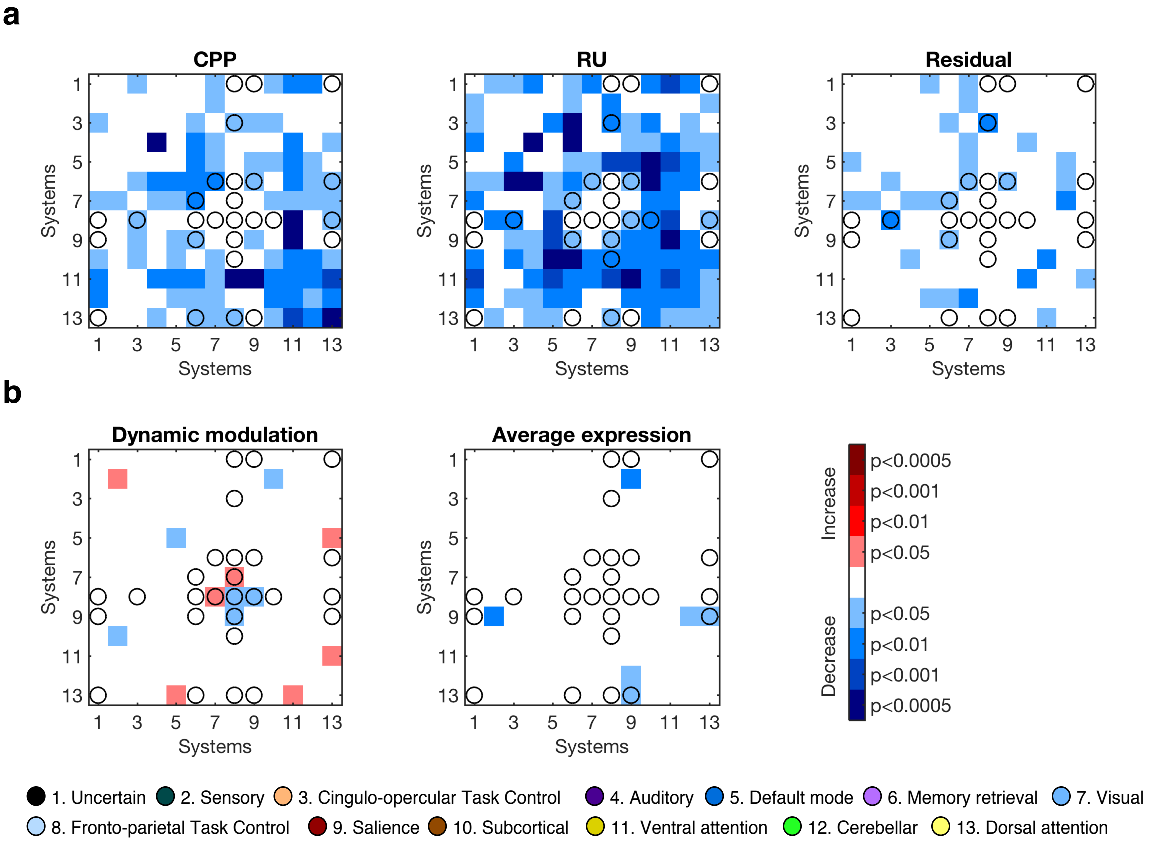


**Supplementary Figure 7**

Contributions of different system edges in subgraph 4. (a) Contributions of different system edges to the effect of learning factors on temporal expression of subgraph 4. We removed all edges for one of the 91 system-by-system connections and re-estimated the coefficients for CPP, RU, reward and residual updating. Then we compared these coefficients with the original coefficients (including all the edges) to estimate the contribution of each system edge. The open circles denote the significant system edges in subgraph 4 (as shown in Fig. 3b). An increase in coefficients is shown in red while a decrease is shown in blue. Lower $p$ values are shown in darker color. A $p$ value around 0.0005 corresponds to a corrected $p$ value of .05 after multiple comparisons (i.e., 0.05/91). (b) Contributions of different system edges to the relationship between normative learning and dynamic modulation and average expression of subgraph 4. We repeated the same procedure and estimated the change of correlation coefficients for each relationship separately.

**a**

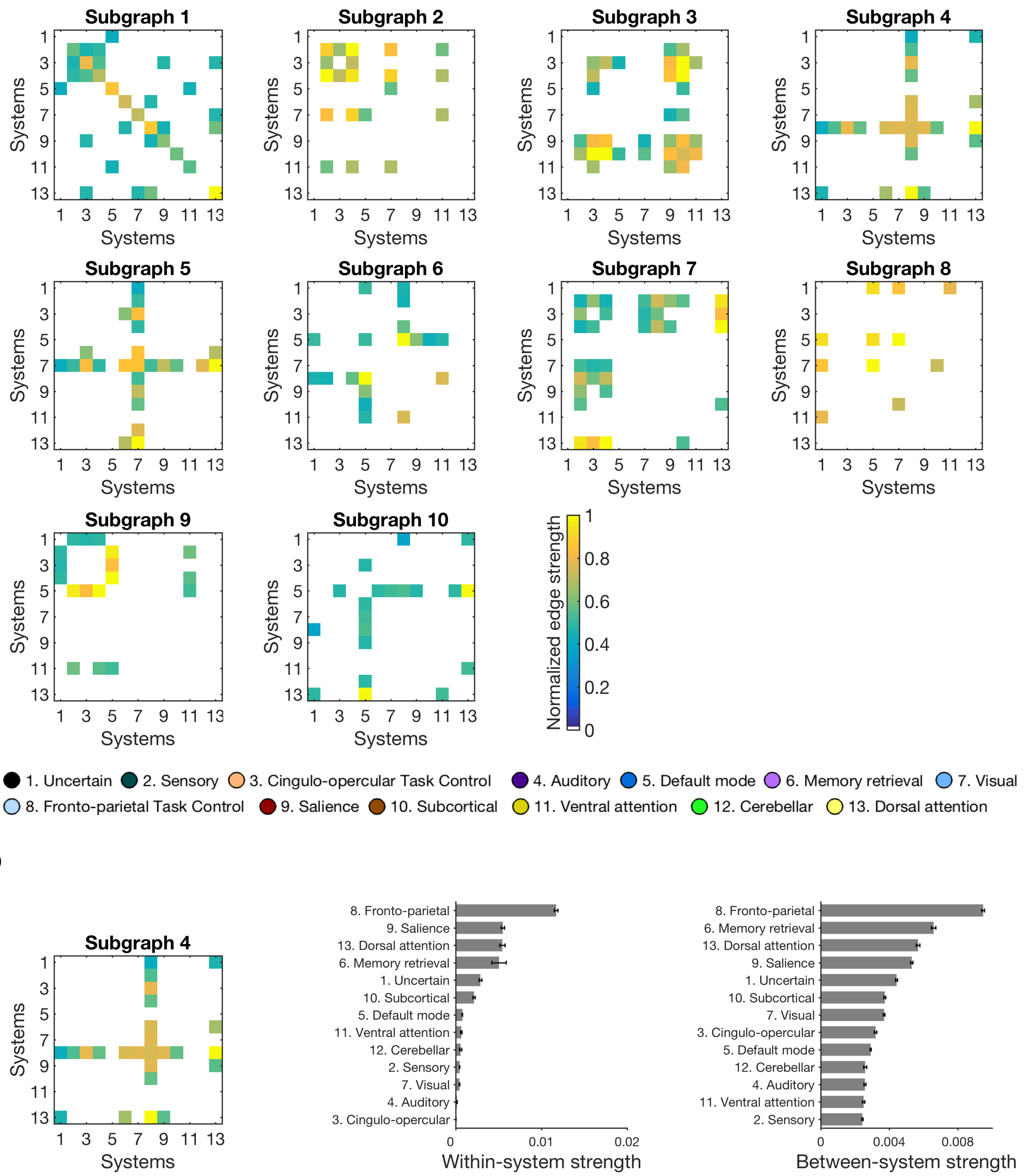


**Supplementary Figure 8**

Robustness check. Subgraphs identified with smaller sliding time window of 8 TRs, with 6 TRs overlapping between time windows. (a) Edges between systems in the ten subgraphs identified by NMF, as in Fig. 3b. (b) Summary of the pattern of connectivity in subgraph 4, as in Fig. 4d, showing the within-system strength and between-system strength of each functional system. The 95% confidence interval of each system was estimated by boostrapping 10,000 times on the edges of that system.

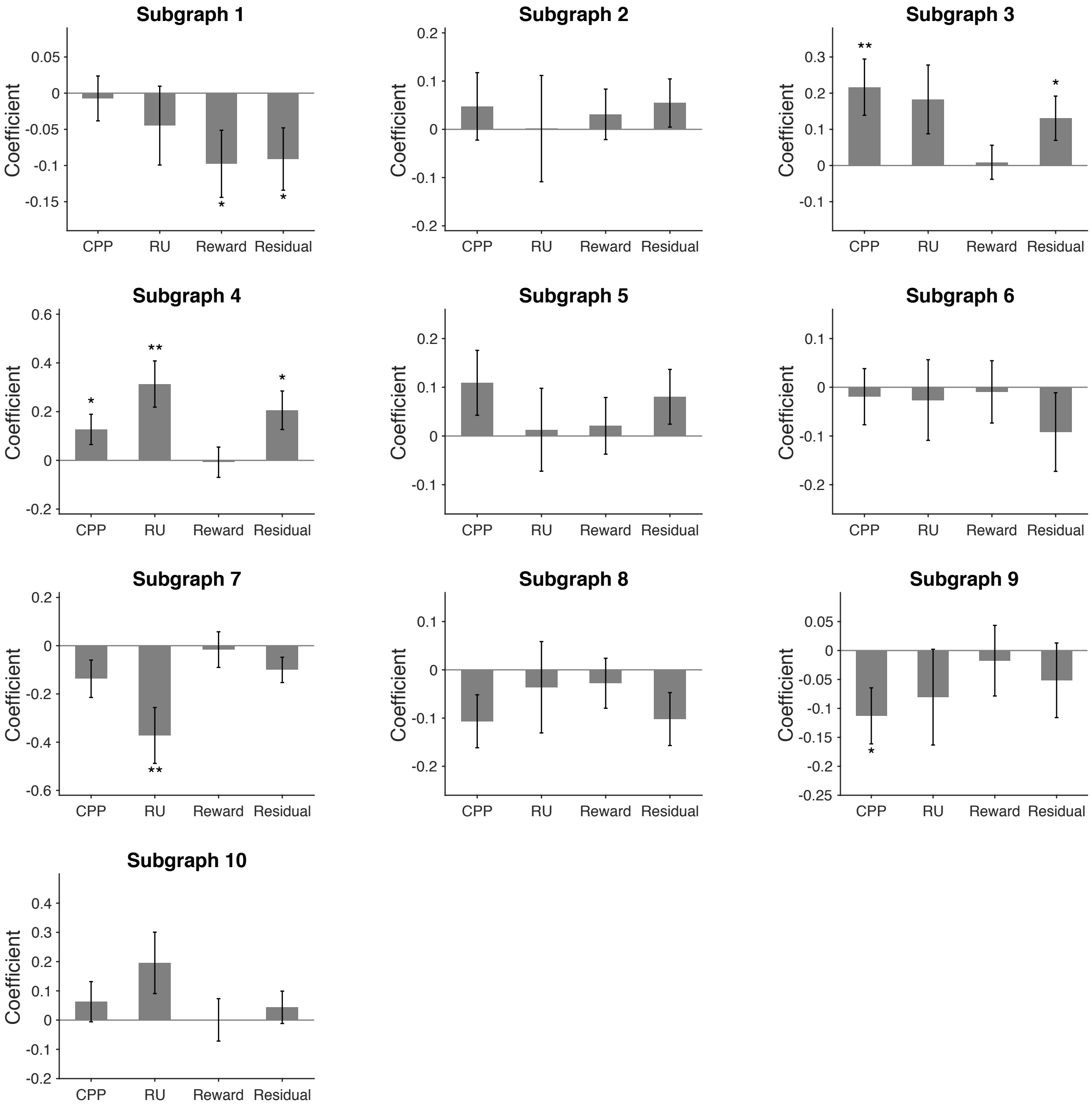


**Supplementary Figure 9**

Robustness check. Modulation of temporal expression by learning factors in all ten subgraphs identified with smaller sliding time window of 8 TRs (Supplementary Fig. 8). Source data are provided as a Source Data file. Error bars represent one SEM. (*$p$<0.05, **$p$<0.01)

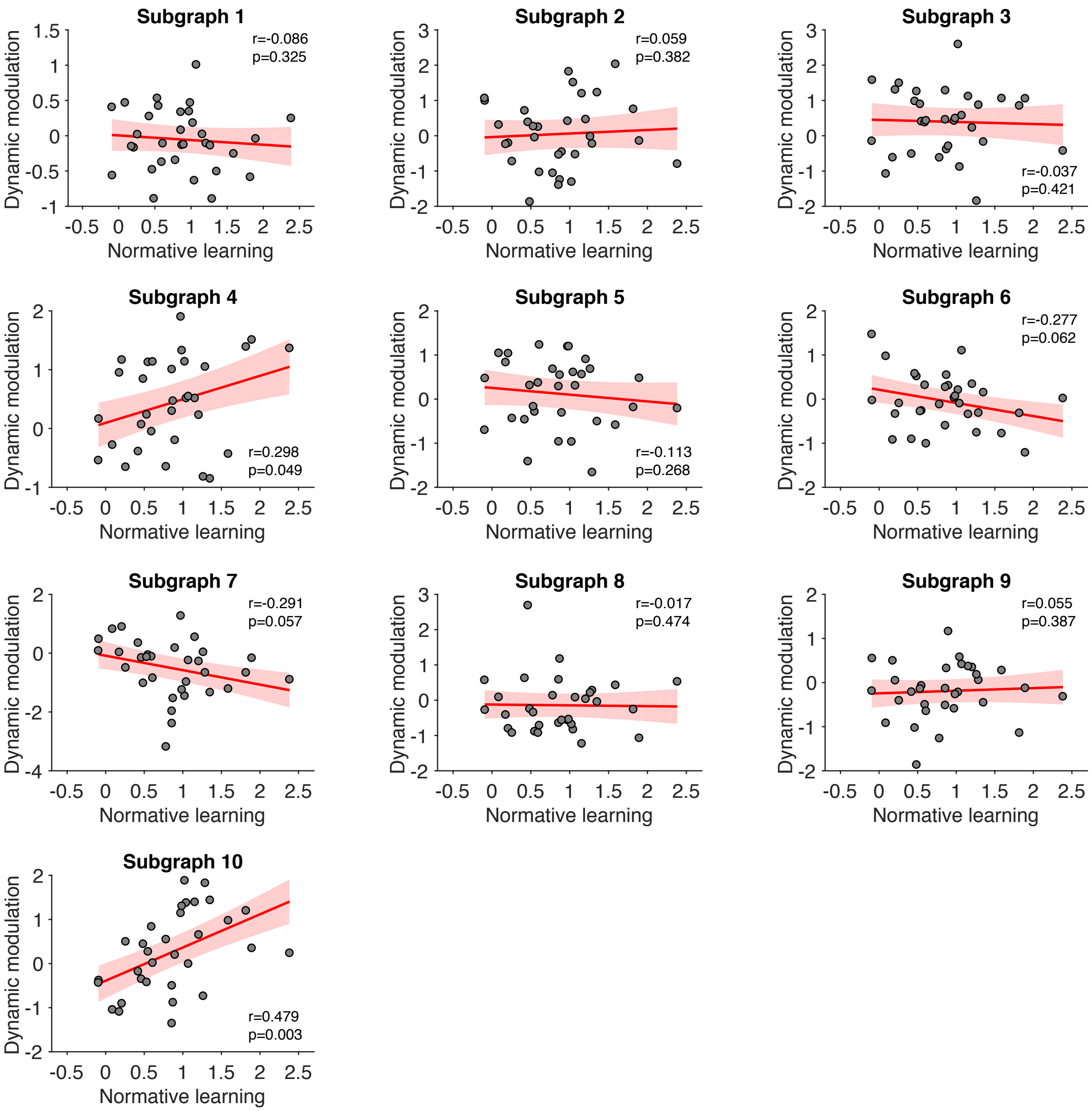


**Supplementary Figure 10**

Robustness check. The relationship between individual normative learning and the dynamic modulation of subgraph expression by normative factors in all ten subgraphs identified with smaller sliding time window of 8 TRs (Supplementary Fig. 8). Each point represents one participant. The red line represents the regression line and the shaded area represents the 95% confidence interval. Source data are provided as a Source Data file.

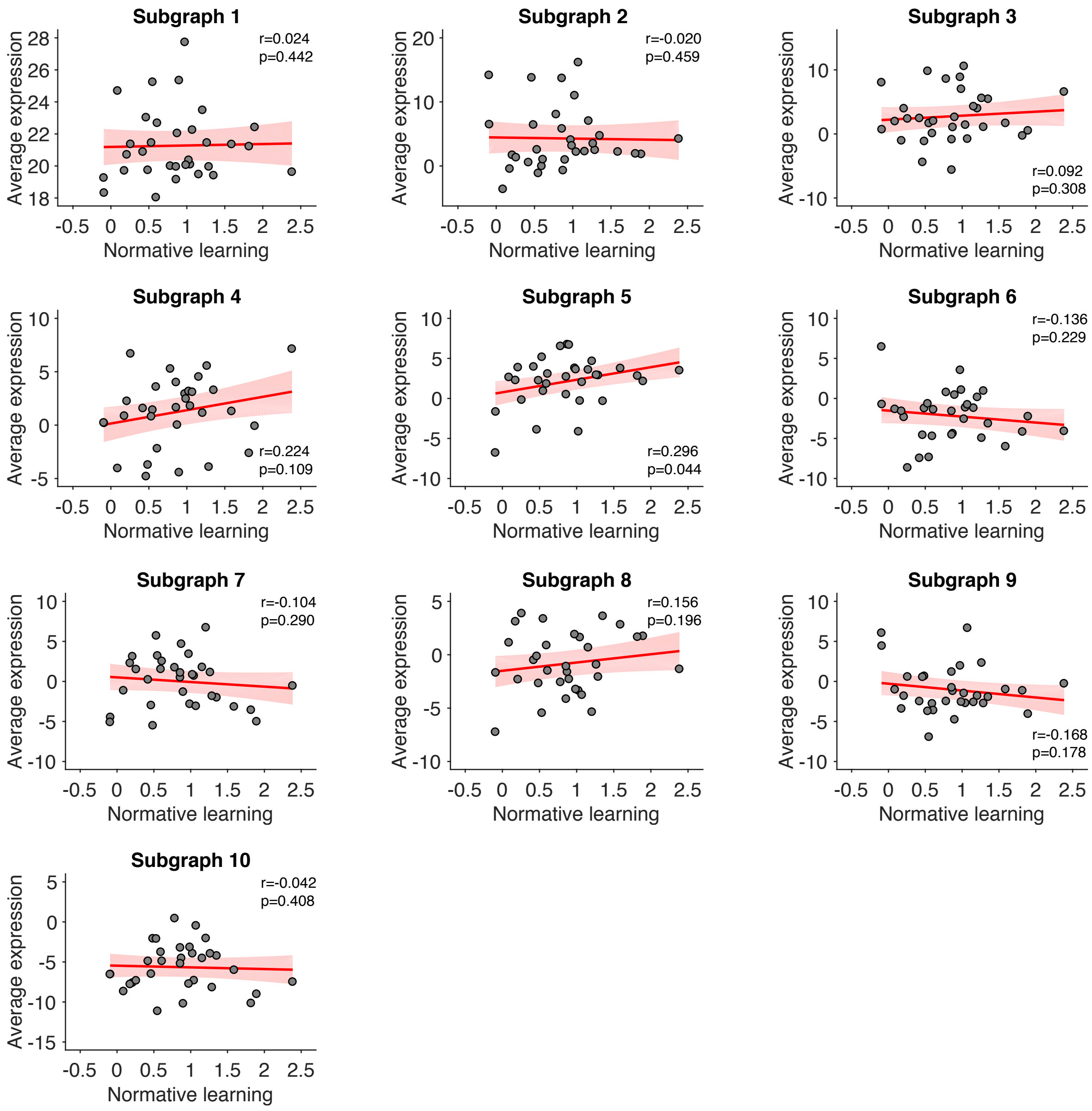


**Supplementary Figure 11**

Robustness check. The relationship between individual normative learning and the average subgraph expression in all ten subgraphs identified with smaller sliding time window of 8 TRs (Supplementary Fig. 8). Each point represents one participant. The red line represents the regression line and the shaded area represents the 95% confidence interval. Source data are provided as a Source Data file.

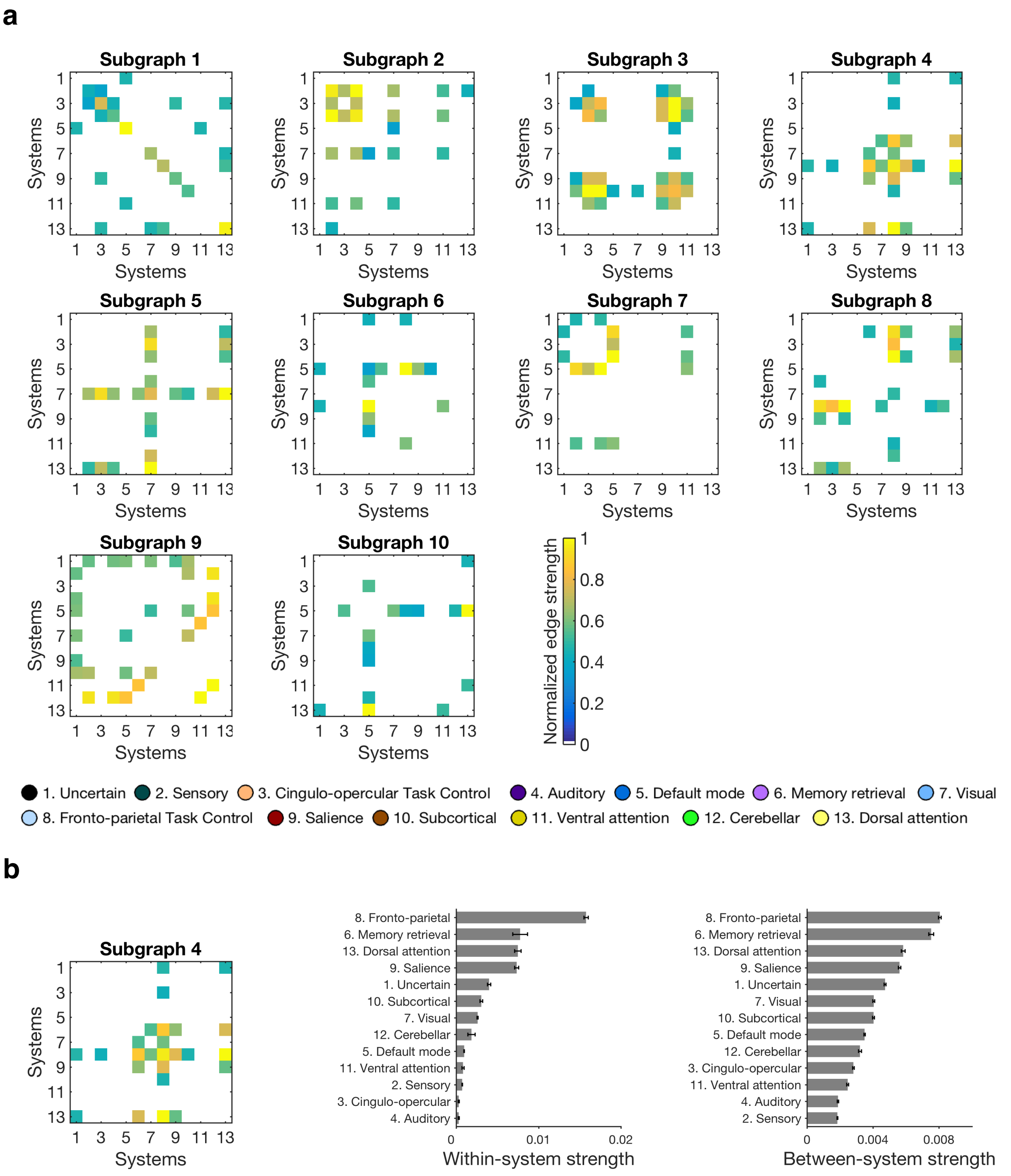


**Supplementary Figure 12**

Robustness check. Subgraphs identified with larger sliding time window of 12 TRs, with 10 TRs overlapping between time windows. (a) Edges between systems in the ten subgraphs identified by NMF, as in Fig. 3b. (b) Summary of the pattern of connectivity in subgraph 4, as in Fig. 4d, showing the within-system strength and between-system strength of each functional system. The 95% confidence interval of each system was estimated by boostrapping 10,000 times on the edges of that system.

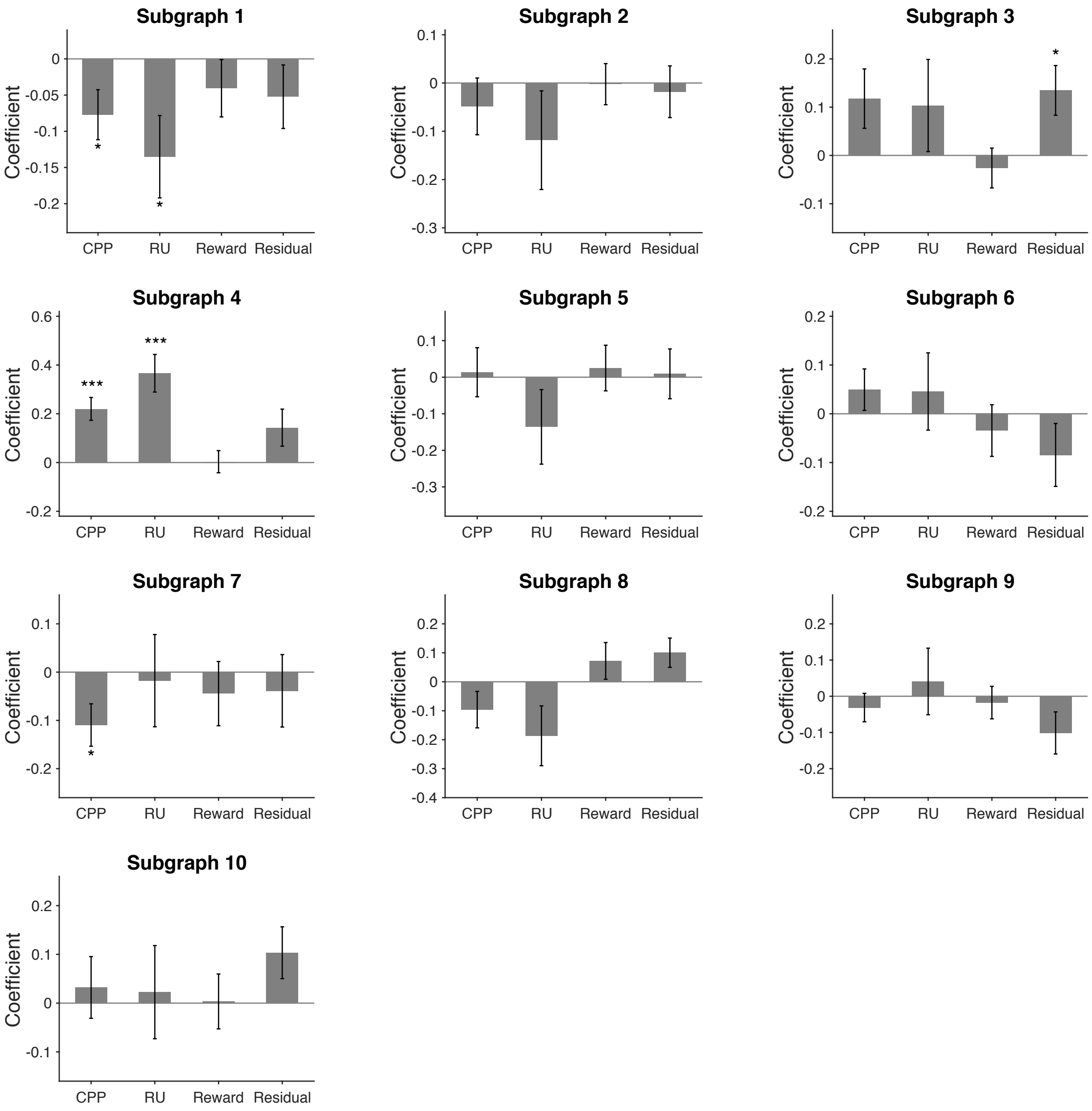


**Supplementary Figure 13**
Robustness check. Modulation of temporal expression by learning factors in all ten subgraphs identified with larger sliding time window of 12 TRs (Supplementary Fig. 12). Source data are provided as a Source Data file. Error bars represent one SEM. (*$p$<0.05, ***$p$<.001)

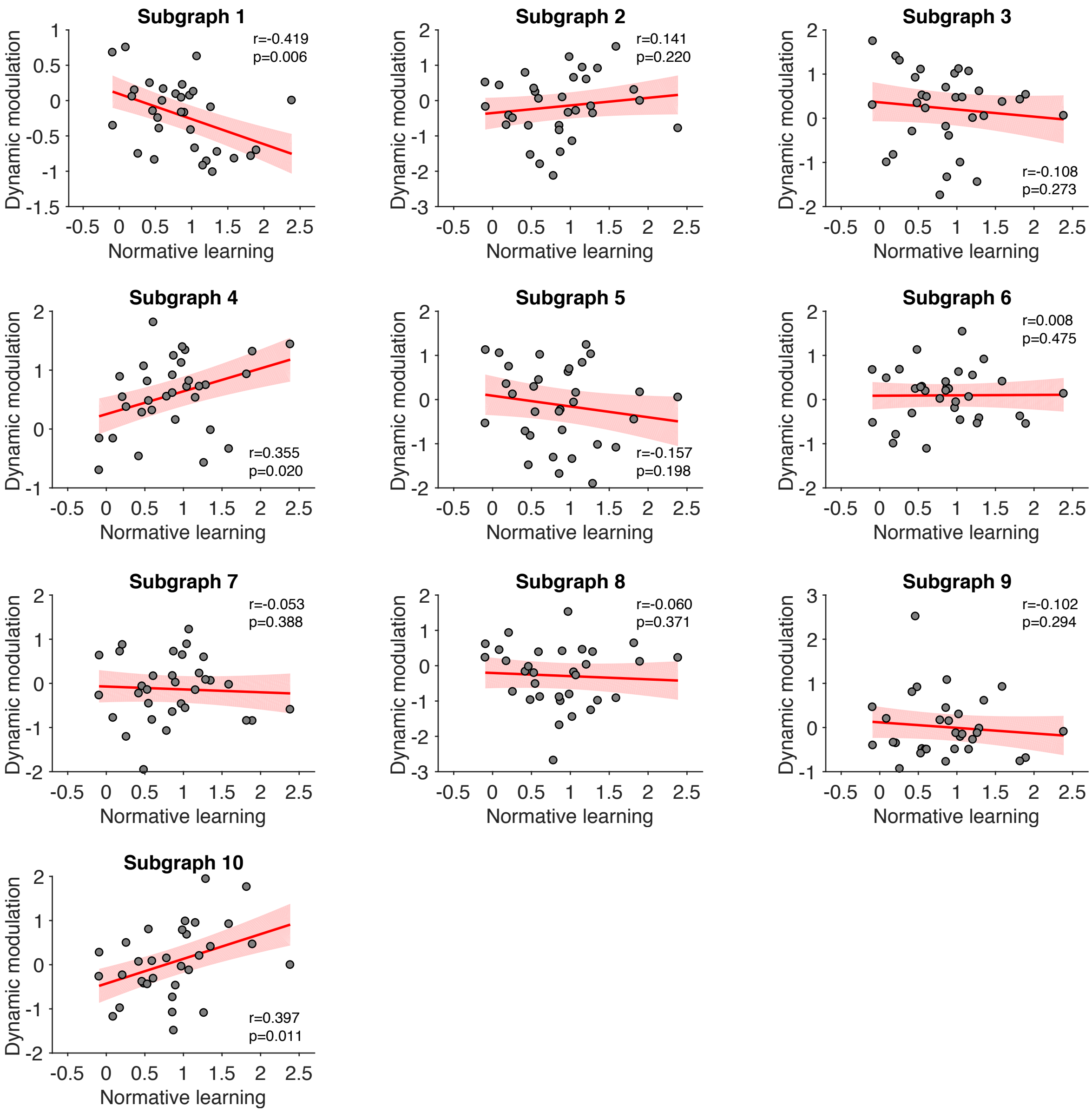


**Supplementary Figure 14**

Robustness check. The relationship between individual normative learning and the dynamic modulation of subgraph expression by normative factors in all ten subgraphs identified with larger sliding time window of 12 TRs (Supplementary Fig. 12). Each point represents one participant. The red line represents the regression line and the shaded area represents the 95% confidence interval. Source data are provided as a Source Data file.

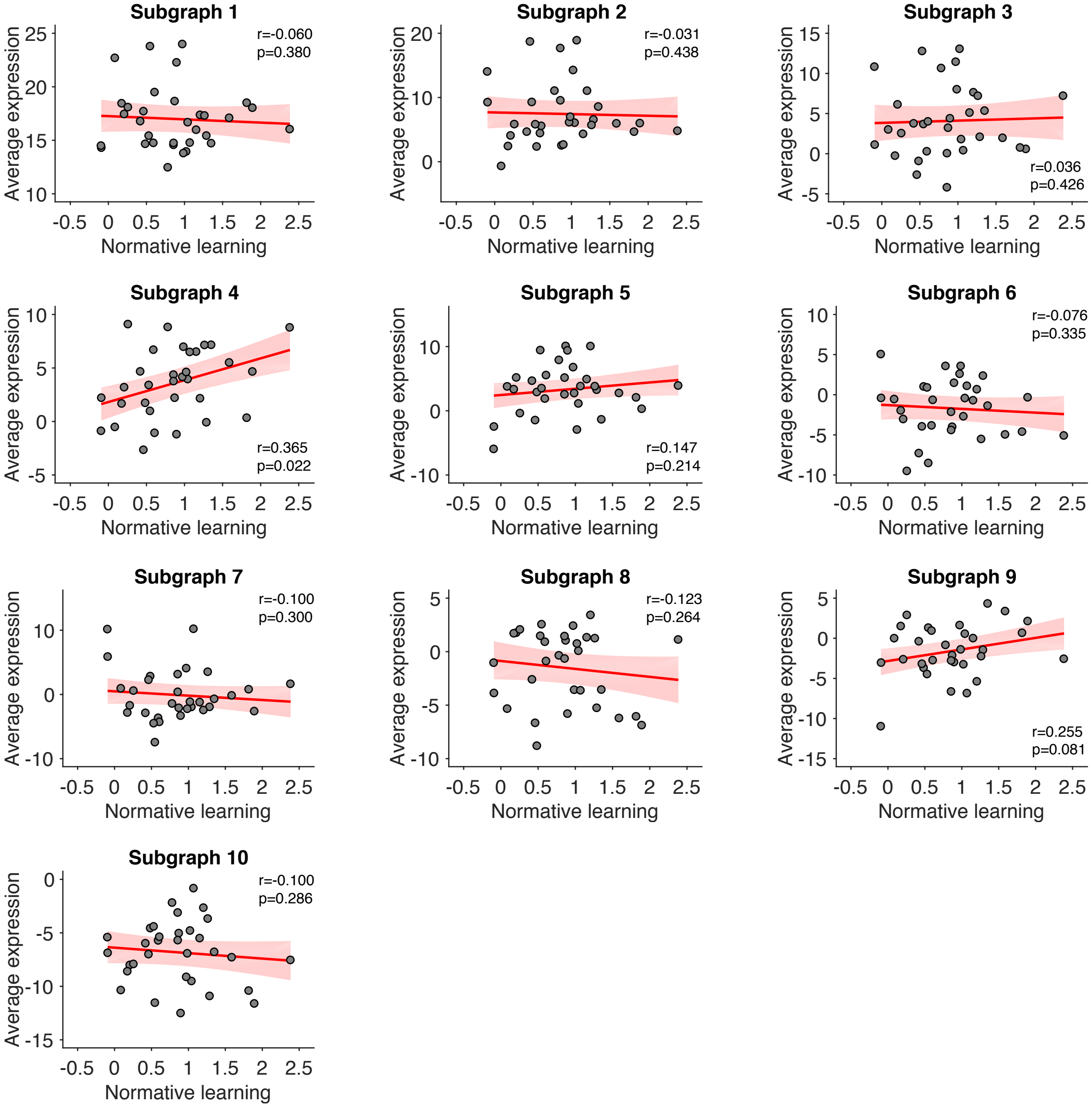


**Supplementary Figure 15**
Robustness check. The relationship between individual normative learning and the average subgraph expression in all ten subgraphs identified with larger sliding time window of 12 TRs (Supplementary Fig. 12). Each point represents one participant. The red line represents the regression line and the shaded area represents the 95% confidence interval. Source data are provided as a Source Data file.

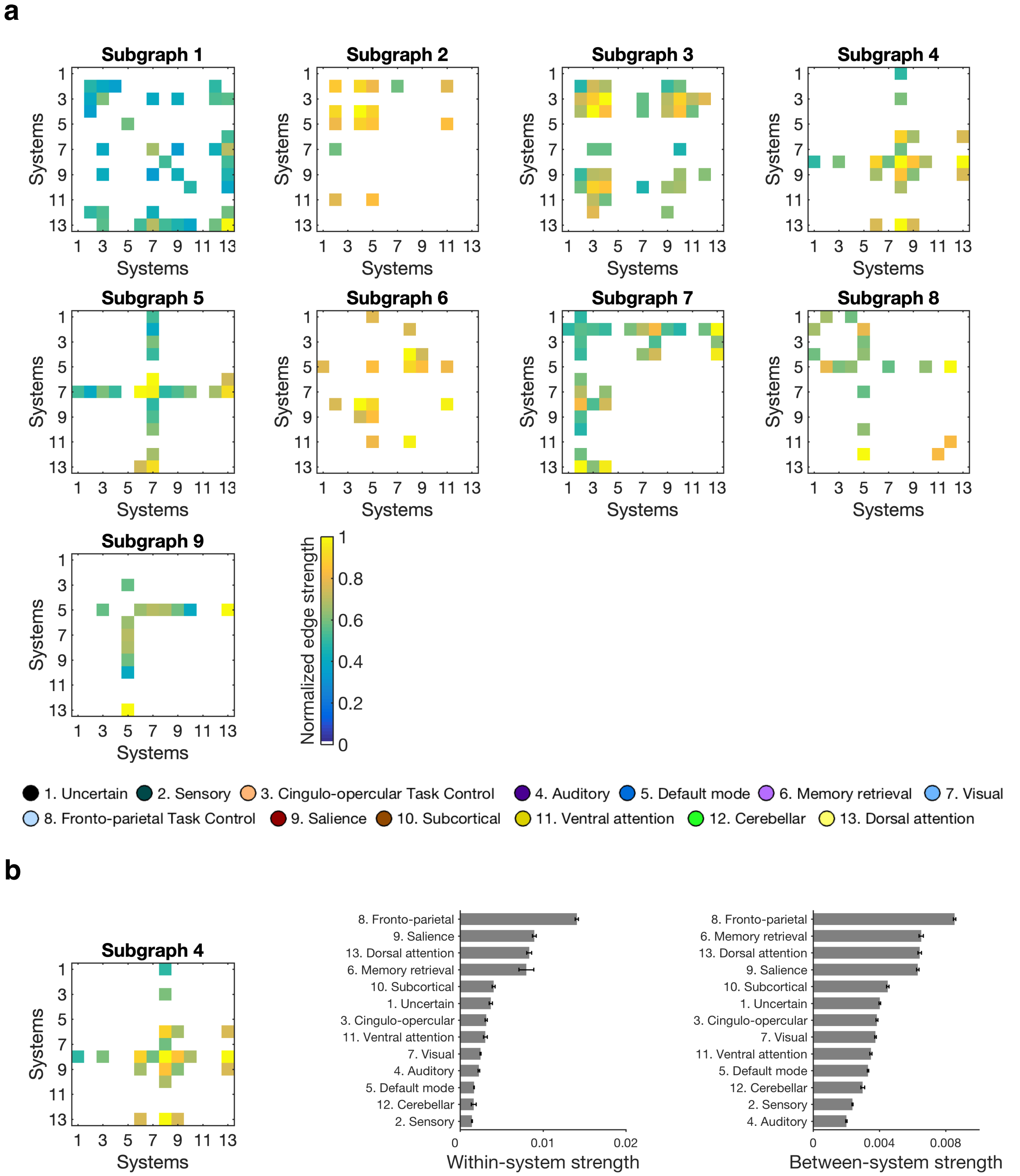


**Supplementary Figure 16**

Robustness check. Subgraphs identified in predicted BOLD signals from univariate GLMs (including predictors for CPP, RU, reward and residual updating). Subgraphs were identified with a sliding time window of 10 TRs, with 8 TRs overlapping between time windows. (a) Edges between systems in the nine subgraphs identified by NMF, as in Fig. 3b. (b) Summary of the pattern of connectivity in subgraph 4, as in Fig. 4d, showing the within-system strength and between-system strength of each functional system. The 95% confidence interval of each system was estimated by boostrapping 10,000 times on the edges of that system.

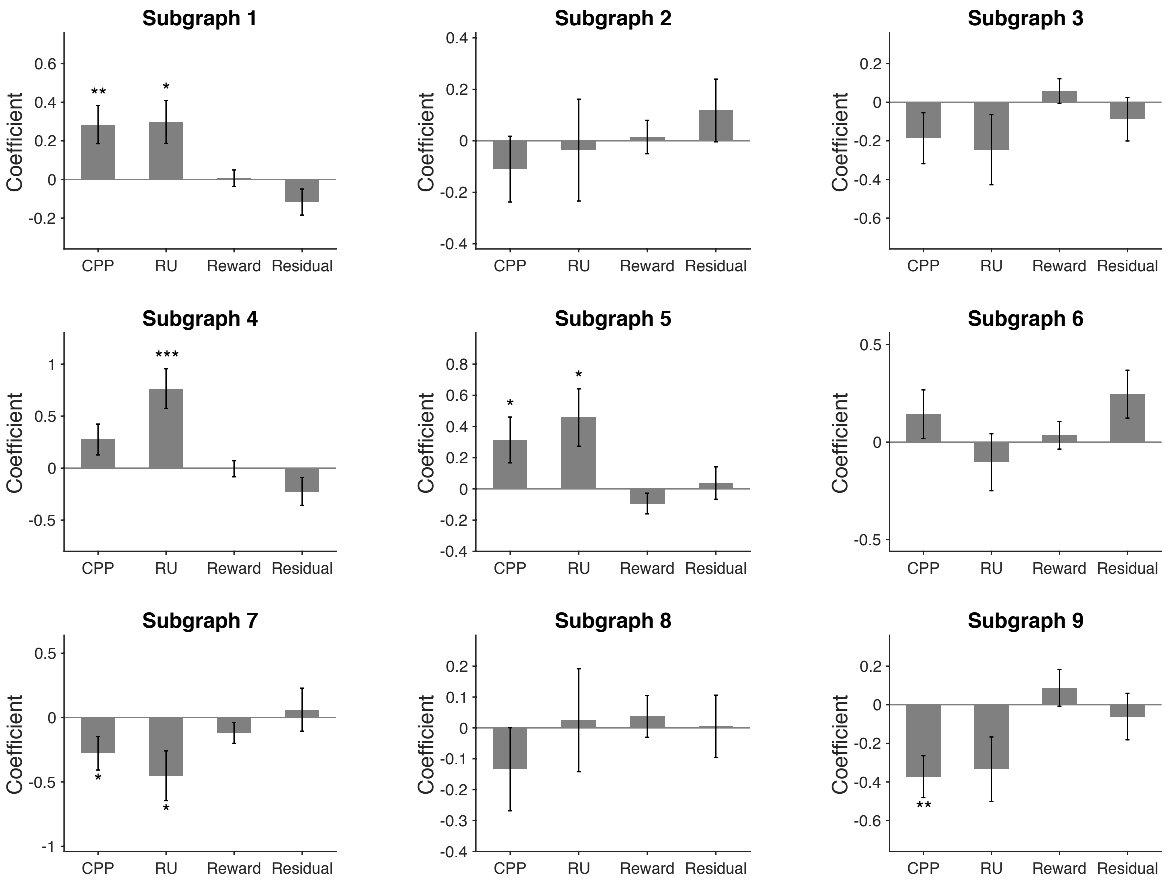


**Supplementary Figure 17**

Robustness check. Modulation of temporal expression by learning factors in all nine subgraphs identified in predicted BOLD signals (Supplementary Fig. 16). Source data are provided as a Source Data file. Error bars represent one SEM. (**p*<0.05, ***p*<0.01, ****p*<.001)

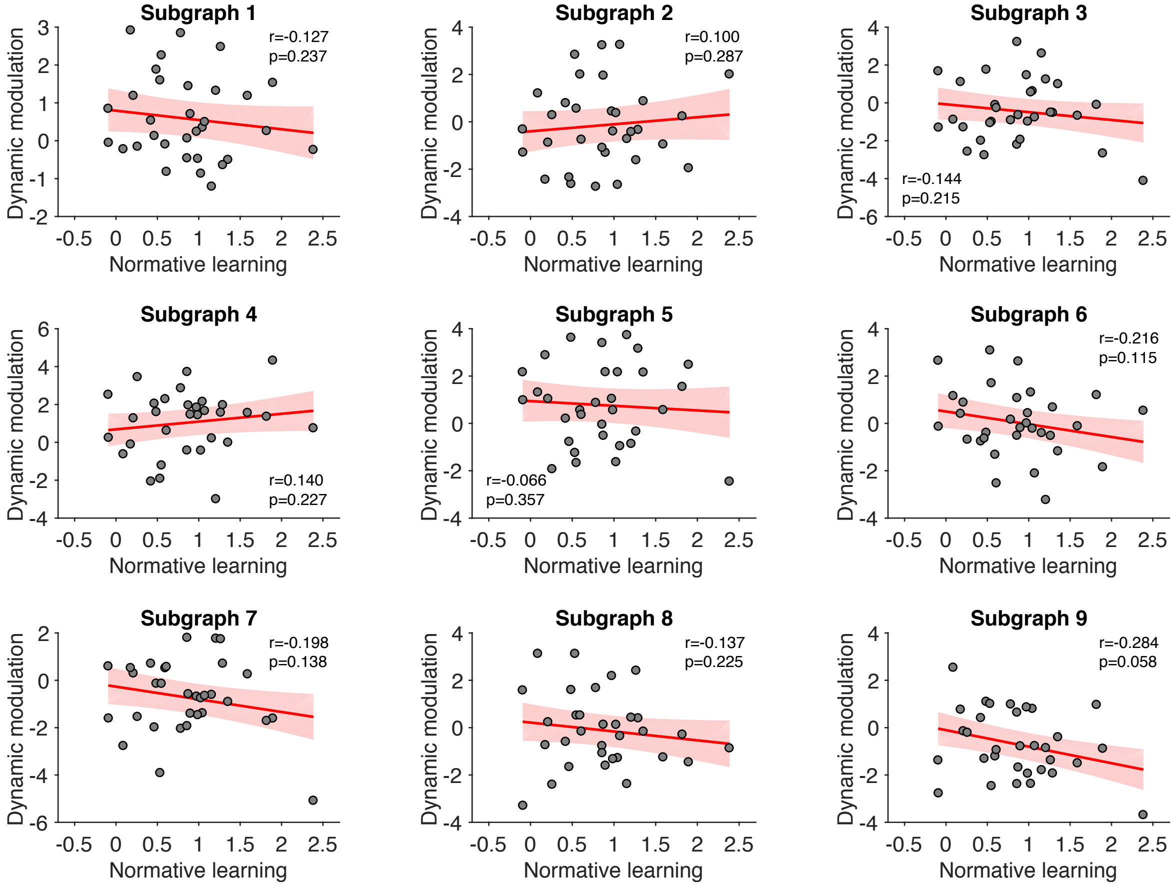


**Supplementary Figure 18**

Robustness check. The relationship between individual normative learning and the dynamic modulation of subgraph expression by normative factors in all nine subgraphs identified in predicted BOLD signals (Supplementary Fig. 16). Each point represents one participant. The red line represents the regression line and the shaded area represents the 95% confidence interval. Source data are provided as a Source Data file.

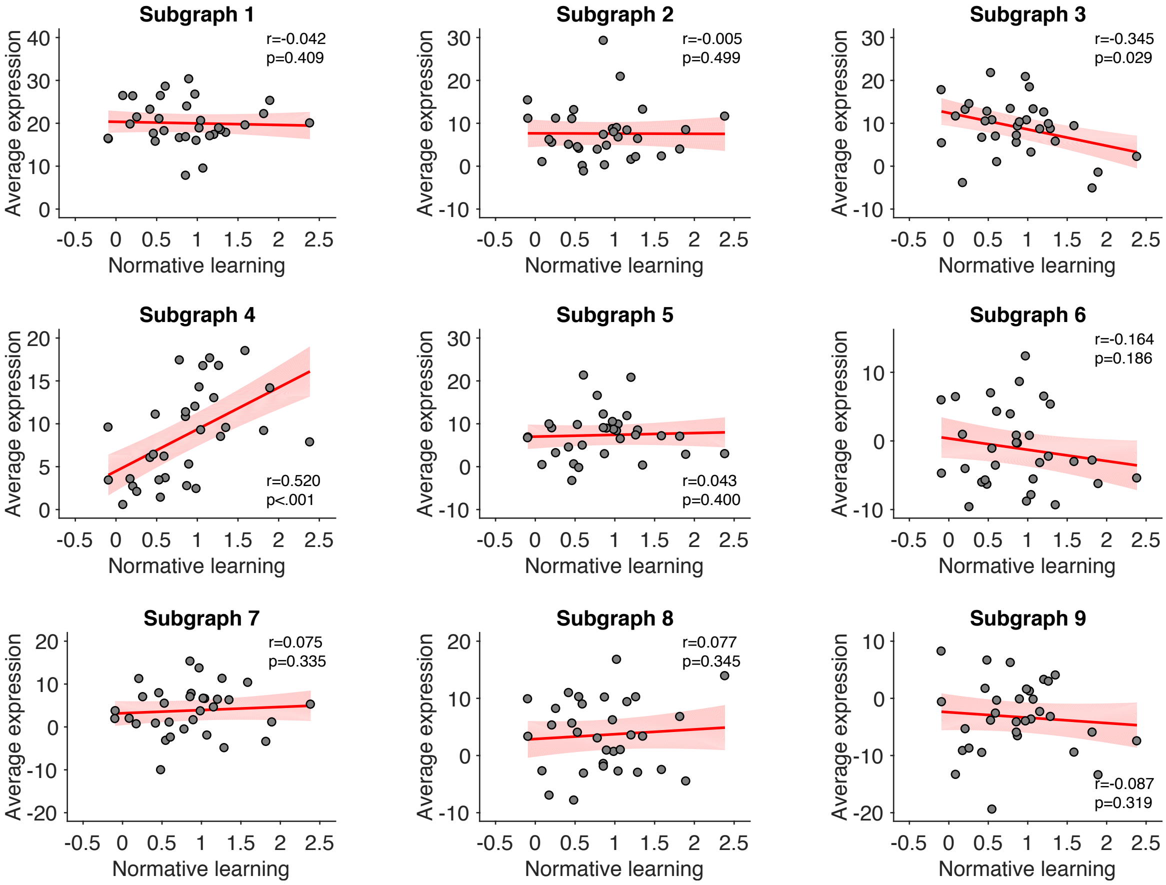


**Supplementary Figure 19**

Robustness check. The relationship between individual normative learning and the average subgraph expression in all nine subgraphs identified in predicted BOLD signals (Supplementary Fig. 16). Each point represents one participant. The red line represents the regression line and the shaded area represents the 95% confidence interval. Source data are provided as a Source Data file.
